# Fabry-Pérot Microscopy for Improved Contrast Enhancement and 3D Cellular Imaging

**DOI:** 10.1101/2025.06.26.661870

**Authors:** Johannes Pittrich, Georg von Köller, Christoph Dillitzer, Daniel Sandner, Ellen Emken, Julia Sistermanns, Zsuzsanna Wolf, Martin Schlegel, Gregor Weirich, Reinhard Kienberger, Oliver Hayden, Hristo Iglev

## Abstract

Fabry-Pérot Microscopy (FPM) integrates a lab-on-a-chip optical cavity and tunable light source in a label-free imaging technique that enhances contrast and enables pseudo-three-dimensional imaging in transparent biological samples. By integrating a fixed-length Fabry-Pérot cavity into a coated microfluidic cell, FPM selectively highlights structures of defined optical thickness through resonance-based interference. We demonstrate that this architecture preserves lateral resolution while providing up to 20-fold contrast enhancement compared to conventional wide-field microscopy. Using human epithelial cells, blood components, and E. coli bacteria, we show that FPM enables clear visualization of subcellular features and discrimination of cell types. Spectrally resolved image stacks are used to extract pixel-wise optical thickness maps, from which physical thickness and refractive index can be derived. These parameters reveal nanoscale structural differences and offer routes to biophysical characterization. Notably, the system operates without mechanical scanning the cavity, using spectral tuning alone to generate images. FPM is compatible with standard microscope optics, and functions under static or flow-compatible conditions, making it suitable for high-throughput cytometry and in vitro diagnostics. These results establish FPM as a versatile extension to wide-field microscopy, enabling contrast-tunable, quantitative imaging of biomedical specimen.

## Introduction

High-resolution optical imaging has undergone intense transformation over the past 30 years, driven largely by innovations in fluorescence microscopy, super-resolution techniques, and optical instrumentation^1–3^. However, despite substantial gains in spatial resolution, limited image contrast remains a recurring difficulty for many biological and clinical applications, particularly when imaging label-free specimens^4–6^. Addressing this contrast limitation is critical for enhancing image interpretability and enabling reliable feature extraction in fields ranging from hematology to in vitro diagnostics.

Several methods have been proposed to increase image contrast, including the use of quantum correlations^7–9.^ Nevertheless, these methods often come with high complexity and cost, limiting their practical application. An alternative pathway using classical light is enabling coherent light to interact with a sample multiple times, this way the signal-to-noise ratio (SNR) can be significantly enhanced mimicking the benefits typically associated with quantum systems^10–13^. In this context, multi-pass microscopy techniques, though conceptually promising, have so far required a lot of optical components and careful alignment. These implementations tend to be bulky, environmentally sensitive, and poorly suited for integration into conventional wide-field microscopes (WFMs). To overcome these limitations, we present a novel approach called Fabry-Pérot Microscopy (FPM), a compact, and mechanically stable imaging method that integrates a on-chip Fabry-Pérot cavity in a microfluidic sample chamber. By plating partially reflective coatings in a microfluidic cuvette, FPM enables light to traverse the sample multiple times, enhancing contrast by nearly an order of magnitude compared to conventional WFM, without compromising spatial resolution. This cavity-enhanced design allows for background suppression, resonant signal amplification, and tomographic imaging, while remaining compatible with existing optical microscopy infrastructure.

The underlying principle of FPM capitalizes on coherent interference within the optical cavity. As light resonates between reflective coatings, its interaction with the sample is multiplied, accumulating phase shifts and enhancing sensitivity to refractive index variations and optical thickness. This capability is particularly relevant in biomedical contexts, where subtle changes in cellular morphology or composition serve as key diagnostic markers. For example, the presence of malaria parasites alters the refractive index of red blood cells^14^, and platelet aggregation levels can offer prognostic insights into various hematological conditions^15^. Beyond simple contrast improvement, FPM facilitates 3D visualization of nearly transparent biological structures. By continuously tuning the illumination wavelength, the optical resonance condition can be scanned, effectively mapping the optical thickness of the sample. Combined with physical thickness measurements, achieved via absorption-based methods using spectral markers, this allows for the extraction of intracellular refractive indices with high spatial precision. In proof-of-concept experiments, we demonstrate a height resolution of approximately 40 nm in optical thickness and a Weber contrast enhancement factor of nearly 20x compared to standard WFM. The compact form and low cost of the FPM platform also lend it to automation and high-throughput applications. For instance, it can be easily integrated into imaging flow cytometry systems, supporting label-free discrimination and counting of blood cells or bacteria^16^. Furthermore, its operation does not require specialized optical alignment, making it particularly well suited for point-of-care diagnostics and widespread clinical use. In general, FPM introduces a practical, efficient, and scalable solution to the long-standing challenge of contrast enhancement in biological imaging. By merging principles of multi-pass interferometry with microfluidic integration, it opens new pathways for detailed label-free analysis of cells in both research and clinical settings.

## Results

### Preservation of Spatial Resolution

To evaluate the spatial resolution and axial imaging behavior of the Fabry-Pérot Microscopy (FPM) setup, we imaged iron(III) oxide nanoparticles with diameters of 50 nm, well below the diffraction limit of a standard microscope using a 0.55 numerical aperture (NA) objective. While the system is not optimized for high-resolution imaging due to its use of parallel illumination, these nanoparticles serve as a useful benchmark for assessing whether the introduction of the Fabry-Pérot cavity degrades the resolution of the microscope. We performed comparative measurements using both uncoated and coated sample cells. Radial intensity profiles around the nanoparticles were extracted and plotted as a function of axial distance from the focal plane, following an approach analogous to Through-Focus Scanning Optical Microscopy (TSOM)^17^. In the uncoated reference cell, the through-focus scan revealed a single, well-defined focal plane with a full width at half maximum (FWHM) of 1.21 µm (Fig. 1a, 1d). In the coated Fabry-Pérot cell, we observed additional axial features. These ghost images are offset by approximately 73 µm and 163 µm from the primary focal plane (Fig. 1b-1c) and are consistent with internal reflections between the partially reflective cavity surfaces. Notably, the reflected images exhibited diminished intensity: the first ghost focus showed a 72% drop relative to the primary peak, and the second a further 65% decrease. These reductions are in line with the expected behavior given the ~68% reflectivity of the cavity coating (Fig. 1e). Despite these internal reflections, the main focal peak in the coated cell maintained a FWHM of 1.20 µm (Fig. 1d), essentially matching the uncoated cell’s resolution within experimental uncertainty.

**Fig. 1.**
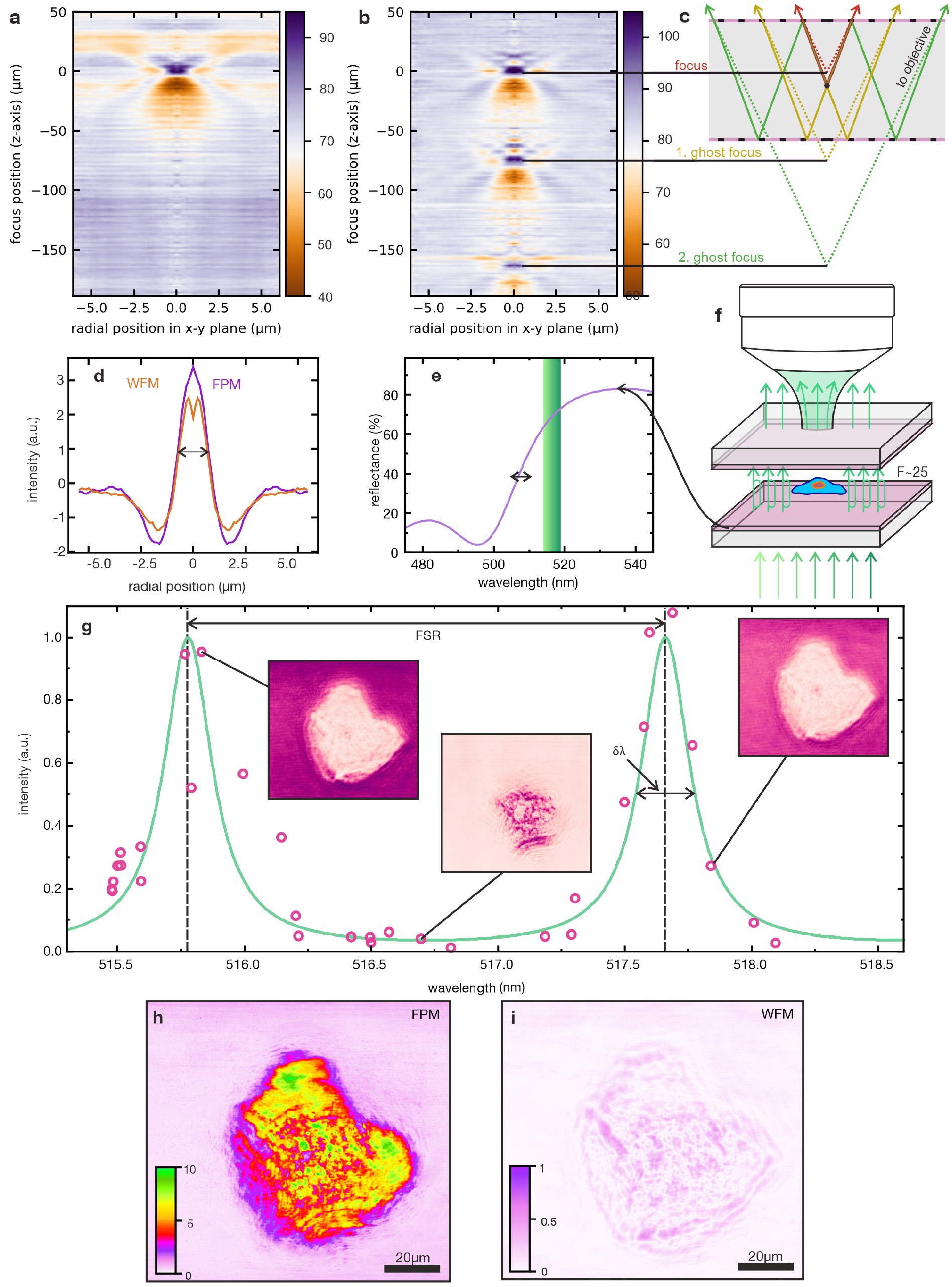
Characterization of FPM. **a**, Radially averaged intensity profile around a 50 nm iron(III) oxide nanoparticle as a function of focus position (z-axis) for the uncoated cell. **b**, Same measurement for the coated FPM cell, showing multiple focal planes due to internal reflections within the cell. Secondary images of the nanoparticle appear approximately 73 µm and 163 µm from the primary focus. **c**, Scheme of internal reflections in the coated cell causing multiple nanoparticle images along the z-axis. Colored rays represent different optical paths contributing to the ghost foci. **d** Point spread function (PSF) of the used microscope in FPM (purple line) and WFM mode (orange) measured with nanoparticles. **e** The reflective coating of the cuvette (purple line) and the tunable laser emission (green). **f** Schematic of multiple light passes through the sample, enhancing the contrast by the Fabry-Pérot finesse factor F. **g** Transmission intensity through the cuvette for different illumination wavelengths (experimental points, calculated green line). Insets: Microscope images at corresponding wavelengths. **h, i**, Weber-contrast image of an epithelial cell measured with FPM and WFM, respectively.

### Contrast Enhancement Through Fabry-Pérot Resonance

A key advantage of FPM is its ability to significantly enhance image contrast by exploiting resonant light transmission through a microcavity (Fig. 1f). To evaluate this enhancement, we used the absolute Weber contrast:

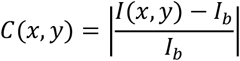

where *I*(*x, y*) is the pixel intensity at a given spatial position and *I*_*b*_ is the local background intensity. This metric is particularly well-suited for imaging small biological structures against homogeneous backgrounds, such as single cells in suspension^18^. To demonstrate FPM’s contrast-enhancing capability, we imaged an epithelial cell at different illumination wavelengths within the cavity’s resonant range. As shown in Fig. 1g, the background intensity adjacent to the cell varied strongly with wavelength. These measured background levels (black points) tracked closely with the theoretical transmission spectrum of the Fabry-Pérot cavity (green curve), confirming that transmission is modulated by the resonant optical thickness of the cavity. Using the above contrast metric, we first established a baseline by averaging over all wavelength within one free spectral range resulting in a standard microscope image. The resulting false-color Weber contrast map (Fig. 1i) revealed a maximum value of 0.55. In contrast, when the same sample was imaged using wavelength-tuned illumination, maximum contrast values increased dramatically across the images. The resulting FPM contrast map (Fig. 1h) revealed local maxima up to 10.78, representing an approximate 20-fold improvement compared to WFM. This enhancement closely matches the cavity’s finesse factor F≈25.

### Optical Thickness Sensitivity and Tomographic Imaging

Beyond contrast enhancement, FPM is sensitive to optical thickness (OT), which allows tomographic information to be extracted from 2D image data. By tuning the illumination wavelength, the system selectively enhances transmission through sample regions where the optical thickness is a multiple of the laser wavelength, enabling mapping of sub-cellular structures based on their refractive index and physical geometry. To quantify this behavior, images were acquired at various illumination wavelengths using a narrowband tunable laser. For each pixel (x,y), the wavelength λ_max_ that produced the highest transmitted intensity was recorded. Using the Fabry-Pérot transmission model, this peak wavelength was mapped to an relative optical thickness value ΔOT(x,y). When applied to epithelial cells, the resulting thickness maps revealed morphological features of the cell body and organelles. Fig. 2a shows a reconstructed image from FPM of an epithelial cell. These features were validated by comparison with a digital holographic microscope (DHM) (Fig. 2b), showing strong agreement in the observed topography. The effective axial resolution of the FPM system was determined by the spectral resolution of the illumination and the order of interference. This results in a resolution limit of Δ*OT*_*limit*_ = 40 nm. By reducing the spectral width of the laser, a resolution limit of around 30 nm could be achieved with the current microfluidic cavity.

**Fig. 2.**
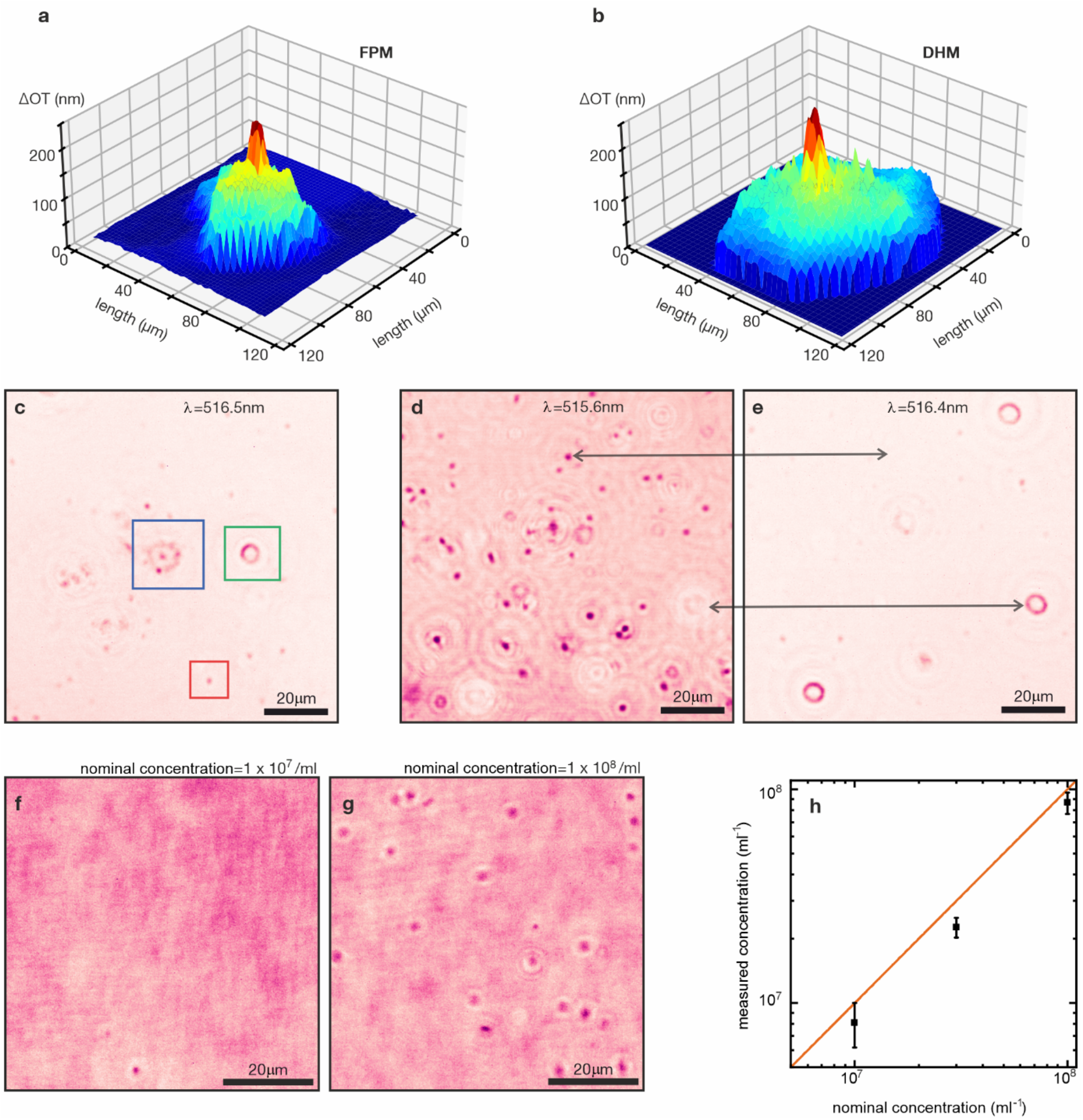
In vitro diagnostic applications of FPM. **a, b**, Tomographic imaging of a human epithelial cell measured with FPM and DHM, respectively. **c**, Different types of blood cells: erythrocytes (green box), immune cells (blue box), and platelets (red box). **d, e**, Thickness-selective imaging of blood sample at two different wavelengths highlighting tailored contrast enhancement. **f, g**, Bacteria measured at two nominal concentrations (nc) of 10^7^ / ml (d) and 10^8^ / ml (e). **h**, Measured versus nominal concentration, while the red line represents the equality of the two values and serves as a guide to the eye.

### Discrimination and Label-Free Identification of Cell Types

Fabry-Pérot Microscopy (FPM) enables label-free differentiation of cell types by using its sensitivity to optical thickness variations, which arise from differences in cellular morphology and refractive index. By tuning the illumination wavelength, specific optical thickness values can be selectively enhanced, allowing visual separation of cell populations within mixed biological samples. To demonstrate this capability, blood samples containing erythrocytes, immune cells, and platelets were imaged using FPM. Fig. 2c shows a representative field of view, where distinct cell classes were identified based on their morphology and contrast. Erythrocytes (green box), immune cells (blue box), and platelets (red box) each exhibited characteristic optical thicknesses, leading to different resonance conditions at specific illumination wavelengths. This wavelength-selective enhancement was further illustrated by imaging the same field at two different wavelengths (Figs. 2d and 2e). Certain cells appeared prominently at one wavelength but were nearly invisible at another, indicating that their optical thickness matched the Fabry-Pérot resonance condition only at specific spectral points. This effect confirms that FPM enables selective visualization of cells based on their individual optical properties, facilitating high-throughput measurement of functional biomarkers at low shear rates while preserving even the mechanically unstable cell aggregates.

### Quantitative Cell Counting and Detection Sensitivity

To evaluate the utility of FPM for quantitative diagnostics, we assessed its performance in detecting and counting small biological objects across varying concentrations. Specifically, we measured suspensions of E. coli bacteria prepared at three nominal concentrations: 10^7^, 3·10^7^ and 10^8^ cells/mL. Imaging was performed under a fixed illumination wavelength using the FPM setup, enabling enhanced detection of low-contrast bacterial structures. Figures 2f and 2g show representative FPM images of bacterial suspensions. At both concentrations, bacterial cells appeared as discrete, high-contrast spots against a homogenous background. To assess quantitative accuracy, the number of detected bacteria in each field of view was compared to the nominal sample concentration. As shown in Fig. 2h, measured counts aligned well with expected values across the tested range. A linear correlation was observed between measured and nominal concentration, with the red line indicating the ideal 1:1 relationship.

### Refractive Index and Physical Thickness Extraction

In addition to imaging optical thickness, FPM enables quantitative extraction of refractive index values by combining its measurements with independently obtained physical thickness data. To demonstrate this, we imaged spherical glass beads suspended in a liquid medium containing an auxiliary dye with selective spectral absorption. The dye was chosen for its minimal absorption at 520 nm, where the FPM operates, and its strong absorption at 740 nm. At the latter wavelength, the dye-filled medium appears dark, while sample regions displace the dye and appear brighter (Fig. 3a). By acquiring images at 740 nm, the physical thickness of the glass beads was determined based on local absorbance contrast, using the Beer-Lambert law. This absorption-based approach yielded a physical thickness of 6.85 ± 0.2 μm, in agreement with the manufacturer’s stated bead size of 7.0 ± 0.15 μm (Fig. 3b, red curve). Using this thickness along with the optical thickness values extracted from FPM measurements, the resulting refractive index of the glass beads was 1.45 ± 0.01 (Fig. 3b, green and blue curves), consistent with known values for the material. These results highlight the potential of FPM as a point-of-care method for rapid analysis pf bacterial infections and antimicrobal resistance.

**Fig. 3.**
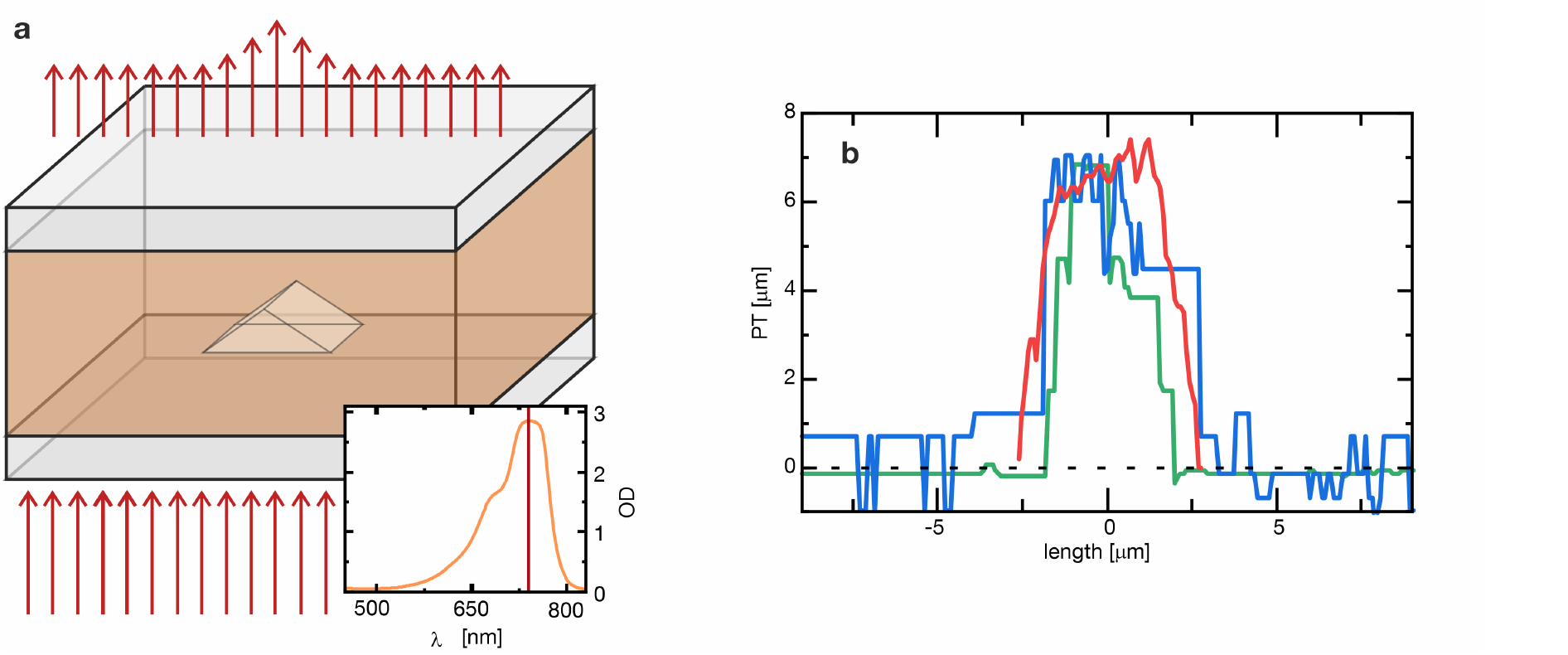
Physical thickness. **a**, Schematic representation of the setup, where light passes through a sample with an additional dye, enabling physical thickness (PT) measurements shown above. Inset: Absorption spectrum of the added dye, with the selected measurement wavelengths indicated in red. **b**, Comparison of physical thickness (PT) profiles of identical glass microspheres obtained through absorption-based measurements (red line) and FPM performed in suspensions with higher (blue) and lower (green) refractive indices of the solvent. The results demonstrate the feasibility of combining FPM with absorption-based techniques to determine both the physical thickness and refractive index of the microspheres.

## Discussion

In this study, we demonstrated that FPM enhances the performance of conventional wide-field microscopy by enabling label-free contrast enhancement, optical thickness-based discrimination, and 3D tomographic imaging. FPM achieved nearly 20-fold contrast improvement without compromising lateral resolution, leveraging a stable optical cavity integrated into a microfluidic cuvette. The technique allowed differentiation of blood cell types and bacterial detection and enabled the extraction of optical and physical parameters such as refractive index and sample thickness. Importantly, these capabilities were delivered with minimal modification to the optical setup and without the need for staining or fluorescence labeling.

The central mechanism behind FPM’s performance lies in resonant enhancement of transmitted light at specific optical path lengths. By tuning the illumination wavelength to match the resonance condition of the Fabry-Pérot cavity, the system selectively amplifies light interacting with structures of matching optical thickness. This provides a powerful method to visualize low-contrast, transparent objects that typically remain undetectable in standard brightfield microscopy. Moreover, FPM’s tunability makes it uniquely suited for spectrally selective imaging. Different cell types or structures, which exhibit small but distinct variations in refractive index and thickness, resonate at different wavelengths. As shown in the discrimination of blood components, this property can be used to highlight specific cell types without chemical labels. The ability to distinguish unlabeled platelets, leukocytes, and erythrocytes in a complex sample represents a significant advancement for label-free cytometry. The technique also enables 3D reconstruction by analyzing wavelength-dependent changes in intensity. Unlike classical tomographic approaches that require mechanical scanning or complex interferometry, FPM uses the spectral information to extract depth information from 2D image stacks. This allows reconstruction of nanometer-scale optical thickness profiles from a compact, stable setup. The observed axial resolution of ~40 nm demonstrates sensitivity comparable to digital holographic microscopy (DHM) while offering simpler integration with conventional optical platforms. Together, these features establish FPM as a versatile imaging method for biomedical research and diagnostics, with potential for applications ranging from in vitro cell analysis to portable point-of-care diagnostic systems.

Compared to conventional WFM, FPM introduces a substantial increase in contrast without altering spatial resolution or requiring sample staining. This positions it as a direct, label-free, drop-in upgrade for existing optical systems. Unlike phase contrast or multi pass imaging, which rely on complex optical path geometry and are limited in spectral flexibility, FPM uses wavelength tuning to achieve contrast selectivity, offering higher adaptability to different sample types and conditions. In comparison to DHM, FPM provides similar depth sensitivity without the need for complex interferometric alignment or phase reconstruction algorithms. The on-chip integration of the Fabry-Pérot cavity within a microfluidic cuvette offers superior mechanical stability, enabling point-of-care deployments. Furthermore, while fluorescence microscopy remains the standard for high-specificity imaging, it often requires labeling steps, photobleaching-prone dyes, and significant sample preparation^19^. FPM, by contrast, is label-free and inherently quantitative, making it better suited for live-cell or longitudinal studies where invasiveness must be minimized. FPM’s compatibility with imaging flow cytometry pipelines also distinguishes it from other multi-pass or tomographic techniques, which often involve bulky or alignment-sensitive optical setups. Its resonance-based mechanism supports both real-time cell identification and count-based diagnostics, which are key needs in clinical hematology, immunology, and microbiology.

Despite its versatility, FPM also presents limitations that must be addressed in future implementations. One inherent constraint is phase wrapping in optical thickness measurements. When the relative optical thickness of a sample exceeds half the illumination wavelength, the system cannot distinguish between consecutive resonance orders, leading to ambiguity unless phase unwrapping is applied. This can be mitigated by dual-wavelength acquisition or extending the spectral tuning range. Another limitation lies in the spectral bandwidth of the light source. The axial resolution and contrast enhancement are fundamentally limited by the laser’s linewidth. While our current system achieves ~40 nm OT resolution with a 0.29 nm bandwidth, commercially available narrow bandwidth sources (bandwith<0.03 nm) could yield further improvements. Similarly, the reflectivity of the cavity surfaces determines the finesse and thus the contrast gain. Increasing mirror reflectivity could enhance performance. Lastly, while ghost foci from internal reflections were not detrimental to lateral imaging, they may introduce complications in dense or thick samples. Careful design of cavity geometry or signal processing approaches could help minimize their impact.

Several opportunities exist to extend and enhance the capabilities of FPM. From a hardware perspective, integrating higher-finesse coatings and tunable light sources with narrower bandwidths could improve both axial resolution and signal-to-noise performance. The use of multiple synchronized wavelengths captured in parallel via dichroic beam splitting or sequence of pulses with different central wavelength could mitigate phase-wrapping issues and increase the accessible dynamic range in optical thickness measurements. Additionally, there is strong potential for combining FPM with other imaging modalities. For instance, coupling FPM with fluorescence microscopy or Raman spectroscopy could combine structural contrast from FPM with vibrational, molecular specificity, enabling multi-dimensional characterization of cells and tissues. Likewise, leveraging the resonance sensitivity of FPM to detect changes in refractive index opens avenues for dynamic imaging of cellular processes such as apoptosis, activation, or osmotic shifts^20,21^. We expect algorithmic improvements to improve robustness and accuracy of obtained tomographic data. From a translational standpoint, the simplicity and robustness of the FPM design make it attractive for integration into compact, automated platforms for clinical diagnostics or environmental monitoring. Its compatibility with microfluidic systems and ability to function label-free may support development of cost-effective point-of-care tools, particularly in resource-limited settings.

## Materials and methods

### Optical Setup and Fabry-Pérot Cuvette

All imaging experiments were performed using a modified wide-field microscope (OKO-1, KERN) adapted for FPM. The system was equipped with a 50× objective lens (NA = 0.55) and integrated with a tunable, narrowband laser source (FPYL-520-02T-TUN, Frankfurt Laser Company), fiber-coupled into the microscope via an added input port. The laser is continuously tunable between 515 and 519 nm, with a spectral bandwidth of approximately 0.29 nm. The tunable laser light was collimated and directed through the sample, which was mounted in a microfluidic Fabry-Pérot cuvette inserted into a custom holder. The transmitted light was collected by the objective and imaged onto a CMOS sensor. All measurements were performed under static conditions (no fluid flow), and image stacks were acquired across the defined wavelength range. To estimate the effective transmission and finesse of the optical system, the laser’s spectrum was cross-correlated with the theoretical Fabry-Pérot transmission function. The system’s measured finesse factor was approximately 20, primarily limited by the spectral bandwidth of the light source.

The FPM cell consists of an industrially fabricated flow cytometry glass cuvette, monolithically etched and coated by IMT Masken und Teilungen AG (Switzerland). The cuvette incorporated a microfluidic channel of several micro meters in height, defined during etching. A partially reflective dielectric coating was deposited on the inner surfaces, forming a stable Fabry-Pérot resonator. The cavity supported resonance-enhanced imaging over the visible spectrum in the 515–560 nm range, where blood components strongly absorb light^22^, with a measured finesse factor up to ~25 at 516 nm. Adhesive bonding into a monolithic structure provided excellent mechanical stability, eliminating the need for realignment or active cavity control. This robust design allowed reproducible imaging conditions and compatibility with standard microscopy hardware. The microfluidic ports enabled sample loading, but for the experiments reported here, static conditions were used during acquisition.

### Sample Preparation

Human epithelial cells were collected non-invasively from saliva and suspended in phosphate-buffered saline (PBS) immediately after extraction. No fixation or staining was applied, and the samples were loaded directly into the Fabry-Pérot cuvette for imaging under static conditions. To preserve their native morphology and optical properties, all measurements were performed shortly after sample preparation. Peripheral human blood served as the source for both erythrocytes and platelets. Platelets were first isolated by centrifugation and then recombined with red blood cells to generate a mixed sample with approximately balanced concentrations. This suspension, likewise unstained and unfixed, was introduced into the cuvette and sealed to maintain a static imaging environment. For reference measurements, silica microspheres (SiO_2_-F-SC31-4, microparticles GmbH) with a nominal diameter of 7.0 µm (±0.15 µm) were suspended in a mixture of ethanol, deionized water, and an infrared-absorbing dye. The dye, 1,1′,3,3,3′,3′-hexamethylindotricarbocyanine iodide (Sigma Aldrich, MKBD8288), exhibits strong absorption at 740 nm while remaining transparent within the FPM spectral window (515–519 nm). This enabled comparative imaging based on absorption contrast. In these measurements, the dye-filled background appeared dark, whereas silica beads displaced the dye and exhibited increased transmittance.

### Data Analysis and Image Processing

Image stacks were acquired by tuning the laser across a spectral range of 515 to 519 nm in 35 discrete steps. Each frame in the stack corresponded to a specific illumination wavelength measured in parallel with a compact spectrometer. For contrast evaluation, the absolute Weber contrast was computed for each pixel at wavelength, then for the same image position, the maximum over all the wavelengths was chosen as the respective maximum Weber contrast and this again for each position. With this, one gets the maximum contrast enhancement independent of the thickness of the underlying sample and therefore easy to compare to WFM. Optical thickness maps were generated by identifying, for each pixel, the illumination wavelength at which the transmitted intensity was maximized. This wavelength, λ_max_, corresponds to the resonance condition of the Fabry-Pérot cavity and is related to the sample’s optical thickness via *OT* = *λ*_*max*_/(2*N*) · *OT*, where *N* is the interference order determined by calibration against background regions. Subtraction of the background reference yielded the relative optical thickness distribution Δ*OT*(*x, y*) = [*n*_*s*_(*x, y*) − *n*_*m*_] ⋅ *PT*_*s*_ (*x, y*). For refractive index extraction, independently measured physical thickness values were used in combination with FPM-derived optical thickness values to solve for *n*_*s*_ via *n*_*s*_ = *n*_*m*_ + Δ*OT*/*PT*_*s*_. Silica microspheres served as calibration standards for this calculation. Their physical thickness was obtained from absorption contrast images using the Beer–Lambert law, while optical thickness was derived from the FPM spectral scan.

